# Non-synaptic plasticity enables memory-dependent local learning

**DOI:** 10.1101/2023.11.14.567001

**Authors:** Ferrand Romain, Baronig Maximilian, Unger Florian, Legenstein Robert

## Abstract

Synaptic plasticity is essential for memory formation and learning in the brain. In addition, recent results indicate that non-synaptic plasticity processes such as the regulation of neural membrane properties contribute to memory formation, its functional role in memory and learning has however remained elusive. Here, we propose that non-synaptic and synaptic plasticity are both essential components to enable memory-dependent processing in neuronal networks. While the former acts on a fast time scale for rapid information storage, the latter shapes network processing on a slower time scale to harness this memory as a functional component. We analyse this concept in a network model where pyramidal neurons regulate their apical trunk excitability in a Hebbian manner. We find that local synaptic plasticity rules can be derived for this model and show that the interplay between this synaptic plasticity and the non-synaptic trunk plasticity enables the model to successfully accommodate memory-dependent processing capabilities in a number of tasks, ranging from simple memory tests to question answering. The model can also explain contextual fear conditioning experiments, where freezing responses could be recovered by optogenetic reactivation of memory engrams under amnesia.

**Author summary:** How memory is organized in the brain in order to enable cognitive processing is a central open question in systems neuroscience. Traditionally, synaptic plasticity is considered the key mechanism for the establishment of memory in the brain. Recently however, this view has been questioned, and it was proposed that non-synaptic plasticity mechanisms play a more prominent role as previously considered. In this article, we propose that both, synaptic and non-synaptic plasticity are central components for the formation and utilization of memory in biological neuronal networks. Our results show that non-synaptic plasticity can act on a fast time-scale to store important information, while synaptic plasticity can adapt network function on a slow time scale in order to facilitate memory-dependent cognitive processing.

## Introduction

Memory is an integral component of biological neuronal systems. It underlies behavior at many levels, starting from basic fear memory to complex cognitive processes such as language understanding. Experimental results have provided ample evidence that memories are stored in so-called memory engrams. The main assumption of the memory engram theory is that learning induces persistent changes in specific brain cells that retain information and are subsequently reactivated upon appropriate retrieval conditions [1–3].

A host of experimental evidence supports the hypothesis that synaptic plasticity is essential for memory storage. However, some recent results indicate that also non-synaptic plasticity such as the regulation of neuronal membrane properties contributes to the creation of memory engrams [4–8]. In fact, there has been some scepticism about the role of synaptic plasticity in memory formation [6, 9, 10]. One important argument is that synaptic plasticity, in particular long-term potentiation (LTP) typically requires repetitive stimulation, whereas learning and memory can arise from single experiences. Another suggestion was that non-synaptic plasticity may act as a permissive signal for synaptic changes [4]. In this view, rapid excitability changes of neurons could set the stage for later synaptic reconfiguration. Taken together, the available studies support the intriguing hypothesis that synaptic and non-synaptic plasticity co-operate to construct memory engrams. However, as stated in [4], the question about the functional role of the plasticity of neuronal membrane characteristics has remained open. In this article, we show that fast changes of the intrinsic excitability in the apical dendritic trunk, which we call trunk strength plasticity (TSP) in the following, can give rise to enhanced learning capabilities of neuronal networks. Our hypothesis is that this non-synaptic TSP, which can rapidly be induced, provides the network with instantaneous memory. This memory is used by the network for memory-dependent computation. Synaptic plasticity on the other hand operates on a slower timescale. The role of synaptic plasticity is thus to learn to make use of the memorization capabilities introduced by TSP for the computational task at hand. In that sense, the two plasticity processes synergize in the search for a task solution (see e.g., [11–13] for other synergistic plasticity approaches).

Our theoretical analysis shows that this division of labor gives rise to temporally and spatially local synaptic plasticity rules. We find that despite the locality of the synaptic and non-synaptic plasticity processes, networks equipped with such mechanisms exhibit remarkable learning capabilities. This is demonstrated for reward-based learning tasks that necessitate instantaneous memory such as a radial maze task as well as for a complex question-answering task [14], thus illustrating that basic language understanding can be acquired through local learning. We finally show that our model is consistent with basic contextual fear conditioning experiments, giving rise to memory engrams that can be reactivated under amnesic conditions, as has been shown experimentally [15].

## Results

### A model for memory processing in pyramidal neurons

Pyramidal neurons are the principal cells of many memory related brain areas such as the hippocampus, amygdala, and prefrontal cortex. They have a characteristic bipolar shape with basal dendrites close to the soma and apical dendrites that are separated from the soma by an extended apical trunk (green schematic cell in Fig. 1A). These two components establish two sites of synaptic integration. They independently integrate synaptic input in a non-linear manner, and these two signals are then combined at the soma to determine the action potential output of the neuron [16, 17].

**Fig 1.**
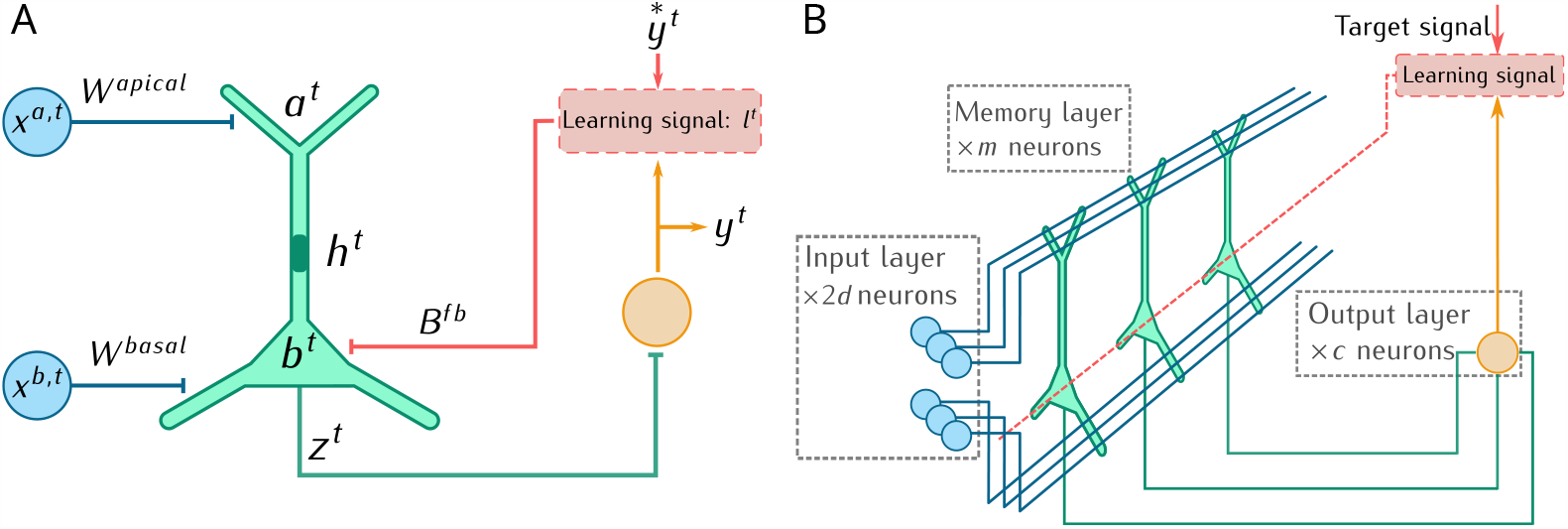
Network model. **(A)** Schematic of our simple pyramidal cell model (green) consisting of an apical and basal compartment with activations *a*^*t*^ and *b*^*t*^ respectively. The excitability of the apical trunk *h*^*t*^ is variable and indicated in dark green. The neuron output *z*^*t*^ is projected to an output layer (yellow). Prediction errors generate learning signals which are fed back via randomly initialized feedback weights *B*^fb^. **(B)** Network view. Input neurons (blue) project to apical and basal compartments of a population of pyramidal cells (green). Pyramidal output **z**^*t*^ is projected to a layer of linear output neurons (yellow) producing the network output **y**^*t*^.

For our network model, we were inspired by memory-augmented neural networks (MANNs) as e.g. in [18, 19]. These models use a memory module where key vectors from ℝ^*m*^ can be associated with value vectors from ℝ^*m*^. When some fact is stored in memory (memorization operation), a key and a value vector is computed and this key-value pair is memorized. When some information is retrieved from memory (memory recall operation), a query vector is computed and a value vector is retrieved based on the similarity between the query vector and all stored key vectors. The retrieved value vector is then used in a single layer neural network to compute the network output. In our model, memories are stored in a *memory layer* that is a population of *m* model pyramidal neurons, see Fig. 1B.

The memory layer receives input from two input populations with activities **x**^a,*t*^ ∈ ℝ^*d*^ and **x**^b,*t*^ ∈ ℝ^*d*^ at time *t*, which project to the apical and basal compartments of the pyramidal neurons via synaptic weight matrices *W* ^apical^ and *W* ^basal^ respectively. In each pyramidal neuron *i* of the memory layer, the resulting basal activation 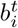 and apical activation 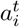 is given by

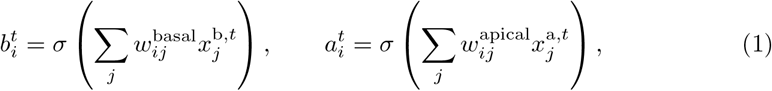

where *σ* denotes a rectified linear nonlinearity, *σ*(*s*) = max {0, *s*} . The vector **a**^*t*^ ∈ ℝ^*m*^ of apical activations of neurons is thus akin to the key vector and the vector **b**^*t*^ ∈ ℝ^*m*^ of the basal activations is akin to the value vector in a MANN. In the following, we will suppress the neuron index *i* in our model description in order to simplify notation. The real valued output *z*^*t*^ of a pyramidal neuron is then given by a linear combination of these activations,

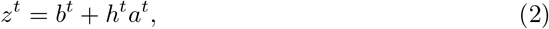

where the scalar *h*^*t*^ *>* 0 denotes the branch strength of the apical trunk. As we will describe below in detail, we model the trunk strength as a dynamic variable *h*^*t*^, used to memorize information about the apical and basal activations (i.e., the key and value vectors). Hence, this layer of pyramidal cells implements a simple memory module. The output **z**^*t*^ ∈ ℝ^*m*^ of the pyramidal cell layer is projected to *c* output neurons (the output layer) via weight matrix *W* ^out^ ∈ ℝ^*c×m*^, which produces the network output **y**^*t*^ ∈ ℝ^*c*^. In addition, the network output can be used to determine learning signals **l**^*t*^ that are used as feedback signals for learning (red in Fig. 1; see below).

A large variety of non-synaptic plasticity mechanisms exist in pyramidal neurons [20–23]. Changes of neuronal membrane characteristics are not necessarily only changing the global neuronal excitability or the action potential initiation threshold, they can also be confined to dendritic subunits [23]. In particular, Losonczy et al. found that local branch activity led to the potentiation of the branch strength when paired either with the cholinergic agonist carbachol or — more importantly — with two or three backpropagating action potentials, indicating an intra-neuron Hebbian type of branch plasticity [23]. Motivated by this finding, we assume a Hebbian-type trunk plasticity:

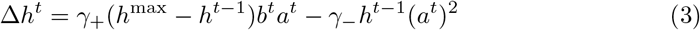

with parameters *γ*_+_ *>* 0, *γ*_*−*_ *>* 0, and *h*^max^ *>* 0 is the maximum trunk strength. Hence, in our model, the trunk strength is potentiated when both the apical and basal compartments are activated by synaptic input (term *b*^*t*^*a*^*t*^ in Eq. (3)). This potentiation saturates at *h*^max^. We have also included a depression term that depends quadratically on the apical activation (last term) similar to Oja’s Hebbian rule [24]. This depression term can reset a memory by depressing trunk strengths of neurons where an apical activation is not paired with a basal one. At an update, we assure that the trunk strength remains non-negative.

From a functional perspective, this plasticity enables the neurons in the memory layer to implement a simple memory system. At each time step *t*, the network can perform either a memorization or a memory recall operation. In a memorization operation, both input populations **x**^a,*t*^ and **x**^b,*t*^ are active and the trunk strength is updated. The layer thus memorizes at which neurons both the apical and basal dendrites were activated. At a memory recall, we assume that only the apical input population **x**^a,*t*^ is active. The activated apical compartments will induce neuronal activity only in those cells in which the trunk strength was potentiated, thus reading out a trace of the memory. The output of the memory layer activates the neurons in the output layer, a learning signal is computed, fed back to the memory layer and synaptic weights are updated according to the plasticity rules described below.

In principle, the apical and basal input populations could provide different aspects of the input at a memorization event. For simplicity however, in our simulations they exhibited the same activity pattern **x**^a,*t*^ = **x**^b,*t*^ at memorization events, which turned out to work well in our simulations.

### Local synaptic plasticity for memory-dependent processing with TSP

In our model, memory is encoded in the network in the vector of trunk strengths **h**^*t*^. This memory is constructed on the fast time scale of single neuronal activations.

According to our hypothesis, synaptic plasticity in contrast works on a slower time scale and is used to learn how to make use of this memory process in the specific task context. Similar ideas have been put forward in a number of models termed memory-augmented neural networks [18, 19, 25, 26]. In these networks however, synaptic weights are optimized with backpropagation through time (BPTT), a complex and biologically highly implausible learning algorithm [27]. In contrast, it turns out that if trunk strengths are used for rapid information storage, one can derive local synaptic plasticity rules, see *Methods*. The derivation takes advantage of the eligibility propagation (e-prop) algorithm [28]. The resulting on-line plasticity rules approximate BPTT using synaptic eligibility traces in combination with learning signals that are directly fed back from the network output to the network neurons. This is illustrated in Fig. 1 for a supervised learning scenario (red). Consider a network with *c* output neurons and corresponding outputs 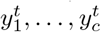. For given target outputs 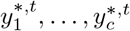, we obtain learning signals 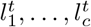 which are fed back to the memory layer neurons through a feedback weight matrix ***B***^fb^ *∈* ℝ^*m×c*^. Consider a single neuron in the memory layer with feedback weight vector ***b***^fb^. The weighted learning signals are summed to obtain the neuron-specific learning signal *L*^*t*^:

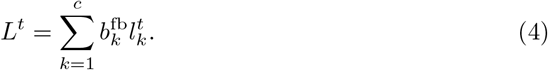

The synaptic plasticity rules for synapses at the basal and apical dendrites are then given by (see *Methods* for a derivation)

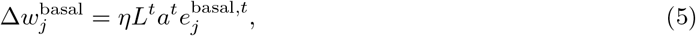

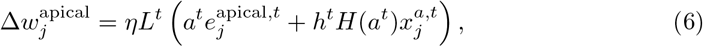

where *η* is a learning rate, 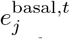 and 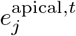 apical synapses respectively, and *H* is the Heaviside step function: *H*(*s*) = 0 for *s* ≤ 0 and *H*(*s*) = 1 otherwise.

In general, the plasticity rules combine the neuron-specific learning signal *L*^*t*^, that is the assigned error to the neuron, with the synapse specific eligibility trace. This eligibility trace records the eligibility of the synapse for changes in the trunk strength. For example, if there was a large error assigned to the neuron and its trunk strength was high, synapses that led to a trunk strength increase are eligible for that error and are changed such that this increase will be smaller in future. The eligibility is computed by filtering information locally available to the neuron:

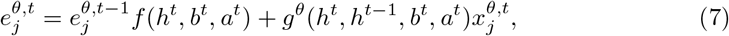

for *θ* ∈ {basal, apical} and functions *f, g*^*θ*^. Note that the update rules (5) and (6) as well as the updates of eligibility traces (7) need only temporally and spatially local information, that is, information that is available at the post-synaptic neuron at the current or previous time step. Hence, this update could in principle be implemented by pyramidal neurons. The function *f* in Eq. (7) dynamically controls the decay of the eligibility, and it is the same for both the apical and the basal synapses:

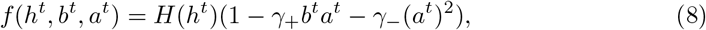

where *H* denotes the Heaviside step function. Hence, *f* (as well as the other functions, see below) are gated by *h*^*t*^, meaning that the eligibility trace is zero if the trunk strength is zero. This arises from the fact that the apical dendrite has no influence on the neuron output when *h*^*t*^ = 0, and thus the network output does not depend on the input to this neuron during a recall (note that during a recall, only the apical inputs are active). It is 1 (no decay) in the absence of apical activity and reduced by the squared apical activity and the product of basal and apical activity. In other words, the synapse stays fully eligible as long as the trunk strength is not altered. If the branch strength is altered due to apical activity, the eligibility is reduced, as other synapses may become eligible for these changes.

The *g*^*θ*^ functions in Eq. (7) modulate the increase of the eligibility trace at synaptic input:

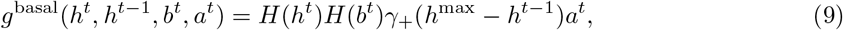

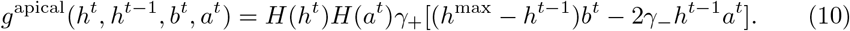

Both are gated by activity in their corresponding compartments. Hence, if the compartment is inactive, the eligibility is not changed even if there is synaptic input, as the compartment did not contribute to any changes of the trunk strength. Also, both include a term *h*^max^ *− h*^*t−*1^ which takes into account that it is harder to increase the trunk strength when it is already close to its maximum value. *g*^basal^ is then linearly dependent on the apical activation, as a potentiation of the trunk strength is proportional to apical activity (Eq (3)). *g*^apical^ is similar, with an additional term that records eligibility according to the decrease of the trunk strength due to apical activation.

The update dynamics of eligibility traces and weights of basal synapses are illustrated in Fig. 2 (in the setup shown there, the apical dynamics are similar). Here, a neuron receives two apical inputs 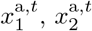 and two basal inputs 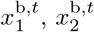 (Fig. 2 right and panel A). First, 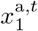 and 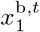 are co-active leading to an increase in the trunk strength (panel B). The eligibility-trace for the first basal synapse is thus increased as it caused this trunk potentiation (panel C). Later, there is a longer co-activation of the other synapse pair 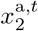 and 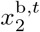, leading to stronger trunk potentiation. Now, the second basal synapse increases its eligibility trace, while the trace of the first one is decreased. At the final time step, a learning signal appears, paired with apical activity. This leads to weight changes that are proporitonal to the eligibility trace (panel E).

**Fig 2.**
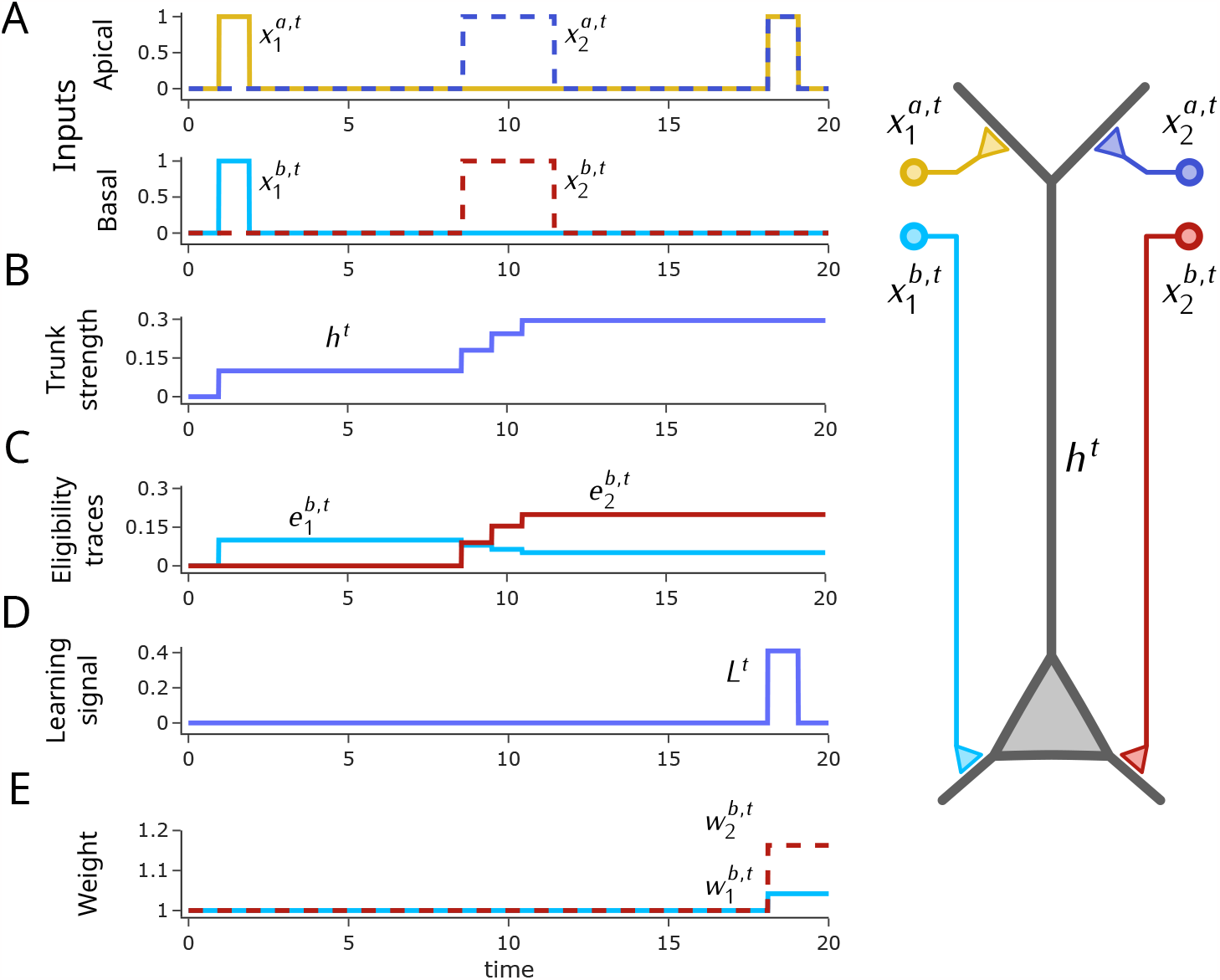
Illustration of eligibility trace dynamics and synaptic plasticity ab the basal compartment in our model. Simulation of a single neuron with two apical and basal synapses, each having unit weight. The first apical and basal synapse is initially activated for one time step, followed by the activation of the second apical and basal synapse for three consecutive time steps (panel A). These co-activations lead to increases in the branch strength *h*^*t*^ (panel B), as well as to changes in the eligibility traces (panel C, see text). Then, a recall is performed at time step 19 where both apical synapses are activated. Further, a learning signal *L*^*t*^ is received (panel D). Changes of basal weights are then given by the product of the eligibility traces with the learning signal and apical activation (panel E, see Eq. (5)). **A)** Inputs to the apical and the basal compartments; **B)** Trunk strength *h*^*t*^; **C)** Eligibility traces for basal synapses; **D)** Learning signal *L*^*t*^; **E)** Basal synaptic weights.

In summary, the eligibility traces record first and second order terms of apical and basal activity, together with some gating and a dependence on the trunk strength.

Although the update rules for the traces are far from simple, they include only terms available at the post-synaptic neuron and in particular are local in time — in contrast to BPTT. Hence, they can in principle be computed at the synapse.

When we compare the synaptic and non-synaptic plasticity mechanism in our model, we observe that they operate on very different time scales with complementary functions. TSP acts very fast in order to memorize relevant information about the current input in the trunk strengths of the memory layer neurons. On the other hand, the local synaptic plasticity dynamics approximate gradient descent and optimize the synaptic weights over many learning episodes in order to minimize network error on the specific task. Their roles are thus complementary, but they synergize in the following way. TSP provides general memory capabilities that help to learn rather arbitrary memory-related tasks. Synaptic plasticity then utilizes this memory and adapts synaptic weights such that the relevant information is memorized and retrieved for the task at hand. In the simulations reported below, we turned off synaptic plasticity in the testing phase after the model was trained. This shows that while the synaptic plasticity is needed for task acquisition, it is not necessary for inference after proper synaptic weights have been determined.

### Learning associations and stimulus comparisons with local synaptic plasticity and TSP

We first tested whether the model is able to learn general associations between sensory input patterns. To this end, we generated two disjunct sets ℛ = {*r*_1_, …, *r*_*n*_}, 𝒮 = {*s*_1_, …, *s*_*n*_}, each consisting of *n* Omniglot [29] characters. The network had *n* output neurons, one for each element in 𝒮 and each element in 𝒮 was associated with one output neuron. Before each episode, we randomly drew a bijective mapping between these two sets to generate *n* stimulus pairs from, ℛ *×*𝒮 such that each element from the two sets appeared in exactly one pair. These pairs were shown to the network in random order, representing facts to be memorized by the network, see Fig. 3A. Afterwards, an input consisting of one element of ℛ was shown as a query and the network had to activate the output neuron associated with the paired stimulus.

**Fig 3.**
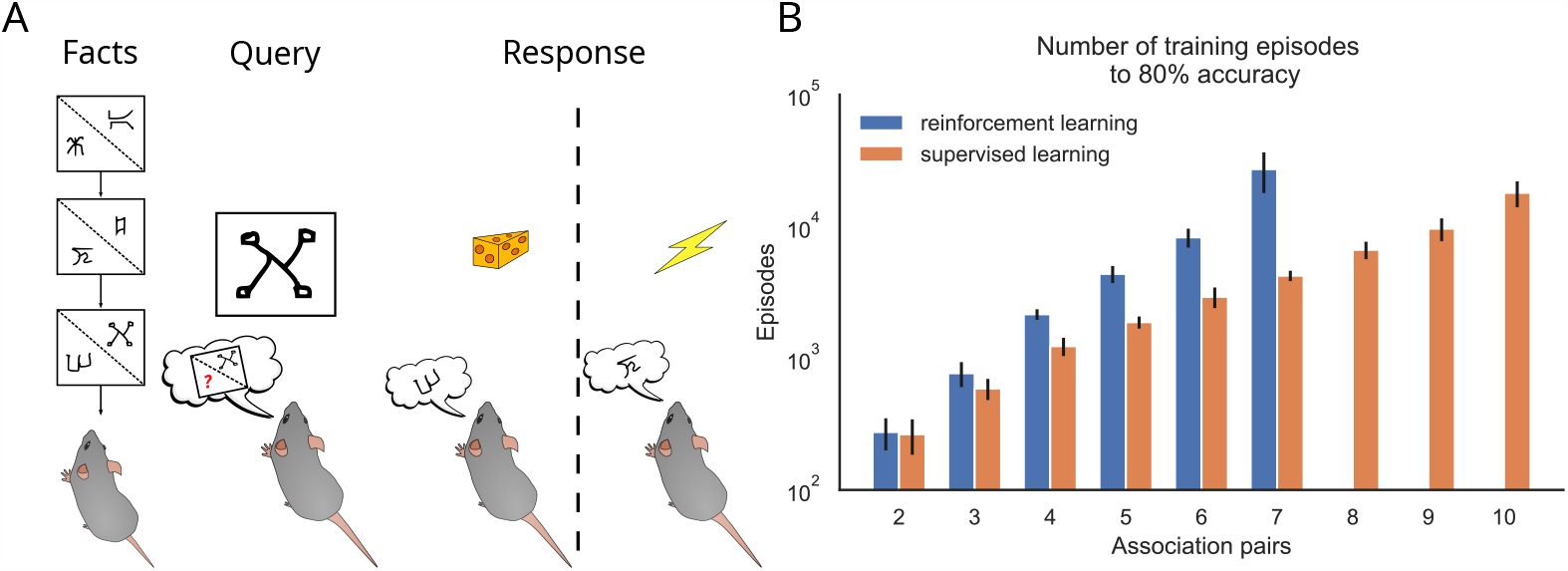
Learning associations with local plastcitiy. **A**) An agent observes a sequence of stimulus pairs. After being cued by one of the observed stimuli, it has to indicate the associated one. **B)** Number of training episodes needed until the network achieved an accuracy of 80% as a function of association pairs to be remembered (mean and SD over 16 training trials).

The network consisted of *d* = 128 input neurons in each of the apical and basal input populations, 200 neurons in the memory layer, and *n* output neurons. In order to generate a reasonable higher-level representation of the Omniglot characters, each character of the presented pair was first embedded in 64-dimensional space with a convolutional network pre-trained using a prototypical objective [30] (the pre-training was done on a subset of the Omniglot classes which were not used in ℛ and 𝒮). The embeddings were then concatenated in a 128-dimensional vector **x**^*t*^. Then, both the apical and basal input populations were activated with this vector, i.e., **x**^a,*t*^ = **x**^*t*^ and **x**^b,*t*^ = **x**^*t*^. At query time, one character from ℛ was embedded and concatenated with a 64-dimensional zero vector to obtain **x**^a,*t*^ to provide the input to the apical compartment, while there was no input to the basal compartment (**x**^b,*t*^ = **0**). We trained the network using a reward-based paradigm where it received a positive reward for the correct response at query time and a negative reward otherwise. In this reward-based setting, we used the standard proximal policy optimization (PPO) objective with an entropy bonus [31] to compute the learning signals used for the synaptic weight updates. The network achieved 100% accuracy on this task after about 2000 episodes for *n* = 5 association pairs. Fig. 3B shows how learning time scales with the number of associations to be learned (blue bars). A few hundred episodes suffice for two associations. For comparison, we also trained the network with direct supervision (error signals are determined from the target response; orange bars). As expected, learning time increase is milder, but with a similar tendency.

Another memory-related task frequently used in experiments is the classical delayed match-to-sample task [32]. Here, the animal observes two stimuli separated in time and must produce an action depending on whether or not the two stimuli are equal, see Fig. 4A. We modeled this task in the setup described above with a pre-trained convolutional network to embed the stimulus in 64-dimensional space and 200 neurons in the memory layer. The agent first observed one out of five Omniglot characters **c**_1_, followed by eight steps where white noise input is shown. Finally, another character **c**_2_ was shown which was chosen to be the same as **c**_1_ with probability 0.5 and one of the other characters with probability 0.125 each. The output of the network was then interpreted as an action ∈ {*a left, righ*}*t* indicating a match or non-match. A reward was delivered accordingly which was used to compute the learning signals for synaptic weight updates. Then, another episode started.

**Fig 4.**
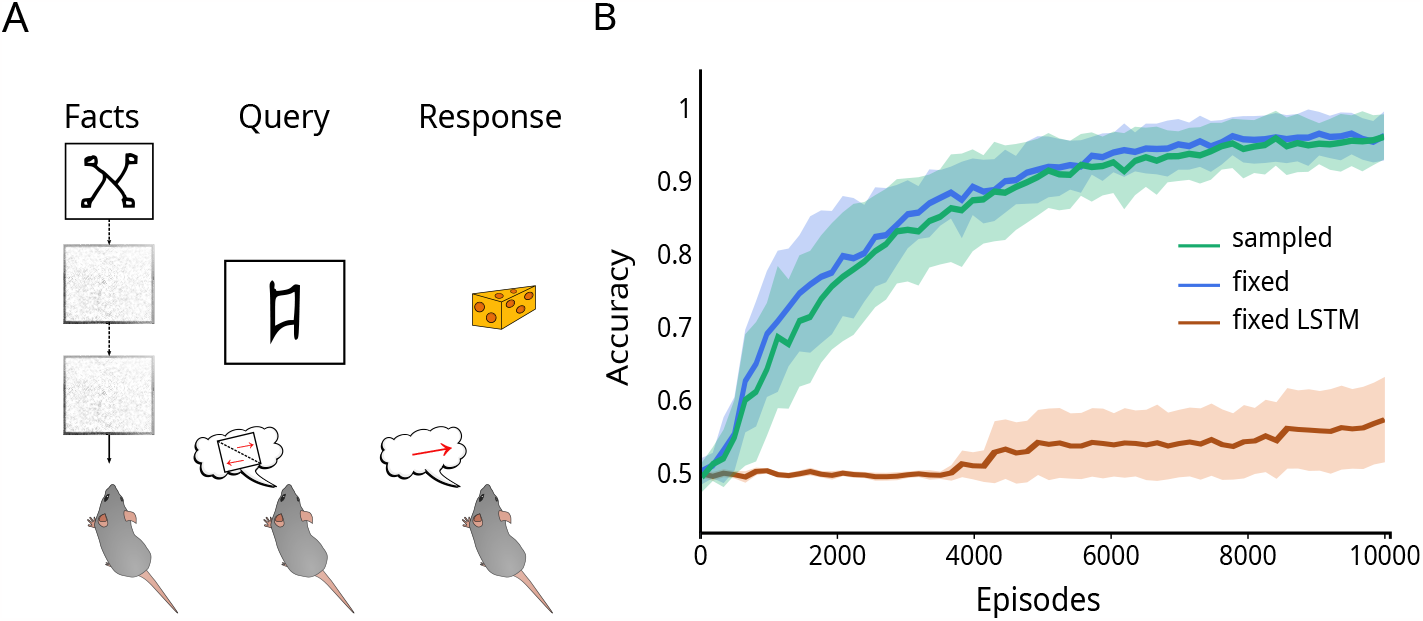
Learning of a delayed match-to-sample task with local plasticity. **A**) Task schema. The agent observes a stimulus **c**_1_ followed by 8 white noise inputs and another stimulus ***s***_2_. The agent should choose the *left* action when the initial stimulus **c**_1_ matches query simulus **c**_2_. **B)** Learning progress in terms of choice accuracy. Green: Only one character instantiation per class for training and testing. Blue: Network is tested on a character not seen during training. Brown: LSTM in the fixed setting (16 trials; shading indicates standard deviation).

In this task, we also tested whether the network can cope with variance in the input stimuli. In Omniglot, each character class consists of 20 drawings of the character from different people with significant variance. We compared network performance in a setting when the specific instantiation of the presented character was fixed to a setting where it was drawn randomly from the set in each episode. In particular, for performance evaluation, we used a character instantiation that was not used for training. Training progress is shown in Fig. 4B. The network achieved an accuracy of 95 *±* 1.8% on this task, with no significant difference between the fixed-character and sampled-character settings. We wondered whether the generalization capabilities of the network in the sampled-character setting could be fully contributed to a writer-independent representation of the characters in the embedding of the convolutional network. We therefore visualized the variance of embeddings for different samples of a given character using principal component analysis and t-SNE [33], see Fig. S1 in the *Supplement*. We found that there is still significant variance in these embeddings. Although the convolutional embedding certainly helps for generalization, this shows that the network can deal with the remaining variance in the input representations.

We also tested the performance of a long short-term memory (LSTM) network [34] trained with BPTT, i.e., with non-local plasticity, see Fig. 4B. Interestingly, the LSTM was not able to learn this task consistently (it reached performances of around 90% in some trials but failed in others). We considered one LSTM with the same number of neurons and one with the same number of parameters as our network. Fig. 4B shows the better performing one. This shows that TSP improves the learning capabilities of neural networks with local synaptic plasticity on this task.

### Learning context-dependent reward associations with local synaptic plasticity and TSP

We next tested whether the network trained with local synaptic plasticity rules was also able to learn a more complex radial maze task [35]. In this task, the animal is located in an eight-armed radial maze (see Fig. 5). It observes one out of four context inputs (in our model, characters from the Omniglot data set), indicating one pair of arms that can be entered (indicated by color in Fig. 5A). One of these arms contains a reward. The animal has to explore the arms and remember the rewarding arm in that context such that, in the next trial within the same context, the rewarding arm can be chosen. Each episode in this task consisted of 40 trials (i.e., 40 context stimuli and arm choices). In each episode, the reward locations were chosen randomly initially and stayed constant throughout that 40-trial episode.

**Fig 5.**
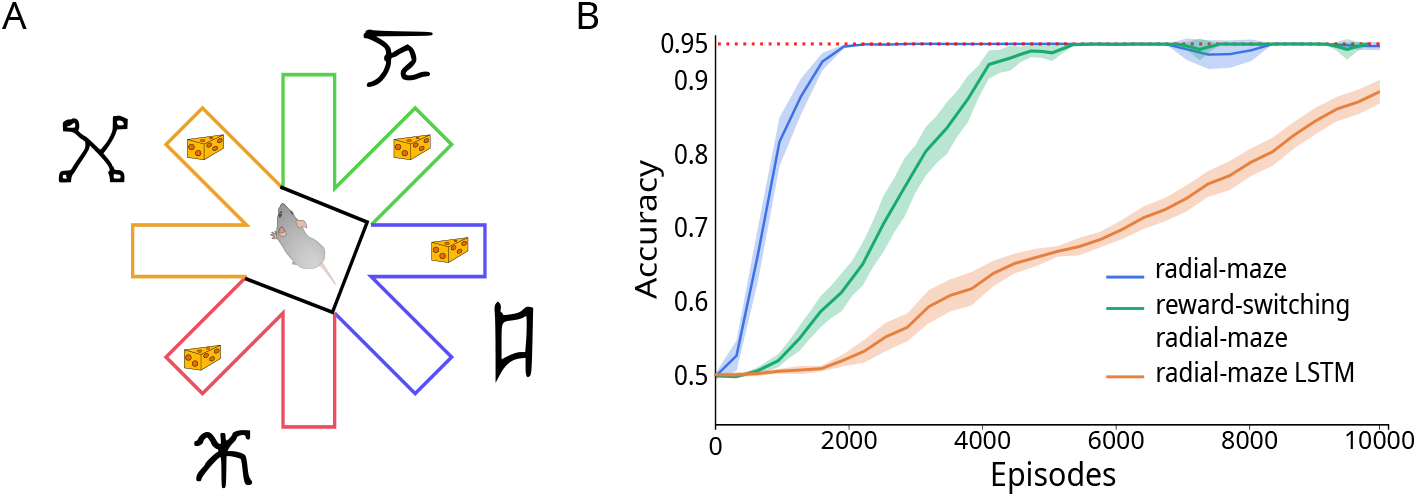
Context-dependent reward associations. **A**) Schema of the radial maze task. In each trial, one arm pair is accessible to the agent (yellow in the example) and the context cue is presented (Omniglot character). The agent then has to choose the correct arm (*left* of *right*) to obtain the reward. **B)** Fraction of rewarded actions over learning episodes in the basic radial maze task (blue) and the same task where the rewarding arm is switched after visit (green; mean and SD. over 16 runs). Red: maximum achievable performance. Orange: LSTM in the basic radial maze task.

We modeled this task using a network as described above, where the visual context was embedded in a 64-dimensional vector using the same pre-trained convolutional network and a 200 neuron memory layer. At the beginning of a trial, one arm pair was chosen randomly out of the four possible pairs and the context stimulus **c** was presented to the network. The network output was interpreted as an action ∈ { *a left, right*} to choose one of the available arms. The network then received either a positive or negative reward, which was used to compute learning signals and update synaptic weights. Afterwards, the network observed a summary of this trial through a triple (**c**, *a, r*), consisting of the context stimulus **c**, the chosen action *a*, and the binary variable *r* ∈ {0, 1 } indicating the received reward. This information could be used by the network to memorize the rewarding action in the given context.

We measured the performance of the network through choice accuracy, that is, the average fraction of rewarding choices within episodes (Fig. 5B). Since the agent has to guess the rewarding action per context initially, the maximum expected accuracy is 0.95. The network learned this task perfectly within about 2000 episodes. Note that this task also includes some basic form of counterfactual reasoning, since the agent can reason about the reward location when visiting a non-rewarded arm. We also tested a more complex variant of this task where the reward in the visited context switches to the other arm after the visit. Also, this task could perfectly be learned with local synaptic plasticity within approximately twice as many episodes compared to the non-switching case. An analysis of the learned network solution can be found in Section 5.2 of the *Supplement*. We also evaluated the performance of an LSTM network with BPTT on the basic version of the task, see Fig. 5B. The LSTM was converging towards a solution, but learned much slower.

### Learning question answering tasks with local synaptic plasticity and TSP

In the above simulations, we tested our model on standard experimental paradigms: a delayed match-to-sample task, and a radial maze task. We next asked whether local synaptic plasticity rules could learn to harness TSP to solve more complex cognitive tasks. One standard benchmark for memory-augmented neural networks is the bAbI task set [14]. It consists of 20 question-answering tasks, where each task is composed of a story consisting of a sequence of up to 325 sentences, followed by a question for which the answer can be inferred from the information contained in the story. See the *Supplement* for example tasks. For our experiments, we used the 10k bAbI dataset to train a network with 200 neurons in the memory layer. According to the benchmark guideline, a task is considered as solved if the error rate is less or equal to 5%. Each sentence of a story was embedded in an 80-dimensional vector, and the sequence of these embeddings was presented to the network sequentially. We first considered a *random embedding*, where we generated an 80-dimensional random vector for each word using the He-uniform variance scaling method [36]. For a given sentence, the vectors of all words in the sentence were then linearly combined with a position encoding that encodes the position of the word in the sentence as in [18].

In Tab.1 we report the mean error rate of the model over 5 runs for each of the 20 bAbI tasks. Using the random embedding, the network was able to learn 13 tasks using local synaptic plasticity (column 3). As a baseline, we considered the same network architecture trained with BPTT (column 2). Notably, all tasks for which the network could be optimized by BPTT could also be learned by our temporally and spatially local synaptic plasticity rules, showing the effectiveness of local learning in our model. Here, the model was trained separately on each of the tasks, resulting in one network for each task. We next tested whether a single network was able to solve all the tasks that could be solved by individual networks by training one model jointly on these tasks (column 4). We found that this was the case and that error rates on a majority of these tasks were even improved, indicating a knowledge transfer between them during learning. In order to test how much a more task-specific sentence representation would improve the results, we also considered a *pre-trained embedding* which is optimized for the task. Here, we used the random embedding as initialization and trained the embedding end-to-end using BPTT on the task at hand. Then the embedding was fixed and a fresh network was trained with local synaptic plasticity on this input representation. We tested all tasks that were not solved with the random embedding, and found that three additional tasks could be solved with a better input representation (column 6). Again, all tasks that could be solved with BPTT could also be learned with local synaptic plasticity.

**Table 1.** Comparison of mean errors of a network trained with BPTT vs our local synaptic plasticity on bAbI tasks 10k (mean and SD. over 5 trials). Error rates for tasks solved by using our local synaptic plasticity rules are printed in bold face. BPTT: backpropagation through time; LSP: Local synaptic plasticiy; LSP joint: joint training where a single network was trained to perform all tasks concurrently.

| Tasks | Random embedding |  |  | Pre-trained embedding |  |
| --- | --- | --- | --- | --- | --- |
|  | BPTT | LSP | LSP joint | BPTT | LSP |
| 1: Single Supporting Fact | 0.3 ± 0.1 | **0.6 ± 0.2** | **0.2 ± 0.1** | | |
| 2: Two Supporting Facts | 55.6 ± 0.9 | 57.8 ± 0.9 | | 35.7 ± 0.4 | 37.4 ± 0.2 |
| 3: Three Supporting Facts | 58.4 ± 0.7 | 62.4 ± 1.1 | | 48.2 ± 0.3 | 48.8 ± 0.5 |
| 4: Two Arg. Relations | 0.02 ± 0.1 | **0.06 ± 0.1** | **0.0 ± 0.0** | | |
| 5: Three Arg. Relations | 1.4 ± 0.2 | **1.8 ± 0.2** | **2.3 ± 0.1** | | |
| 6: Yes/No Questions | 1.0 ± 0.3 | **1.4 ± 0.2** | **2.0 ± 0.1** | | |
| 7: Counting | 1.5 ± 0.2 | **2.6 ± 0.4** | **1.4 ± 0.2** | | |
| 8: Lists/Sets | 1.5 ± 0.3 | **0.9 ± 0.2** | **2.3 ± 0.1** | | |
| 9: Simple Negation | 2.3 ± 0.2 | **2.4 ± 0.2** | **0.3 ± 0.1** | | |
| 10: Indefinite Knowledge | 3.3 ± 0.1 | **3.2 ± 0.1** | **3.2 ± 0.3** | | |
| 11: Basic Coreference | 3.3 ± 0.2 | **3.1 ± 0.2** | **3.0 ± 0.1** | | |
| 12: Conjunction | 2.1 ± 0.2 | **2.7 ± 0.2** | **0.4 ± 0.1** | | |
| 13: Compound Coref. | 7.0 ± 0.5 | 7.5 ± 0.5 | | 2.7 ± 0.1 | **2.9 ± 0.2** |
| 14: Time Reasoning | 15 ± 0.8 | 14.5 ± 0.8 | | 10.1 ± 0.2 | 11.8 ± 0.3 |
| 15: Basic Deduction | 0.6 ± 0.2 | **1.0 ± 0.4** | **0.8 ± 0.3** | | |
| 16: Basic Induction | 48 ± 10 | 51.3 ± 2.1 | | 38.8 ± 3.6 | 47.8 ± 2.4 |
| 17: Positional Reasoning | 19 ± 0.5 | 17.4 ± 1.0 | | 3.7 ± 0.5 | **4.7 ± 0.5** |
| 18: Size Reasoning | 4.7 ± 0.1 | **4.8 ± 0.3** | **2.6 ± 0.4** | | |
| 19: Path Finding | 56.3 ± 0.5 | 61.3 ± 0.9 | | 1.9 ± 0.1 | **2.6 ± 0.3** |
| 20: Agent's Motivations | 0.0 ± 0.0 | **0.0 ± 0.0** | **0.0 ± 0.0** | | |

### Overlapping assembly representations emerge through local learning

How does the network solve such tasks after training? In order to answer this question, we analyzed the behavior of a trained network in the “Single supporting fact” task, see Fig. 6. This task involves simple person-location relations such as “John moved to the kitchen” among several persons and possible locations. Although the results are presented with the verb “moved”, we note that there are actually various phrases in the story to indicate such person-location relations. Those variations do not alter the person-location relation, and the model should thus learn to treat them as equal. To analyze stories, we recorded the vectors of trunk strengths **h**^*t*^ that arose from the dataset’s stories. These vectors thus represent the memory state of the memory layer throughout the stories. Subsequently, we applied a non-negative matrix factorization (NMF) [37] to project these memory states into a 20-dimensional space. We then projected the memory states, keys, values, and recall keys into this space.

**Fig 6.**
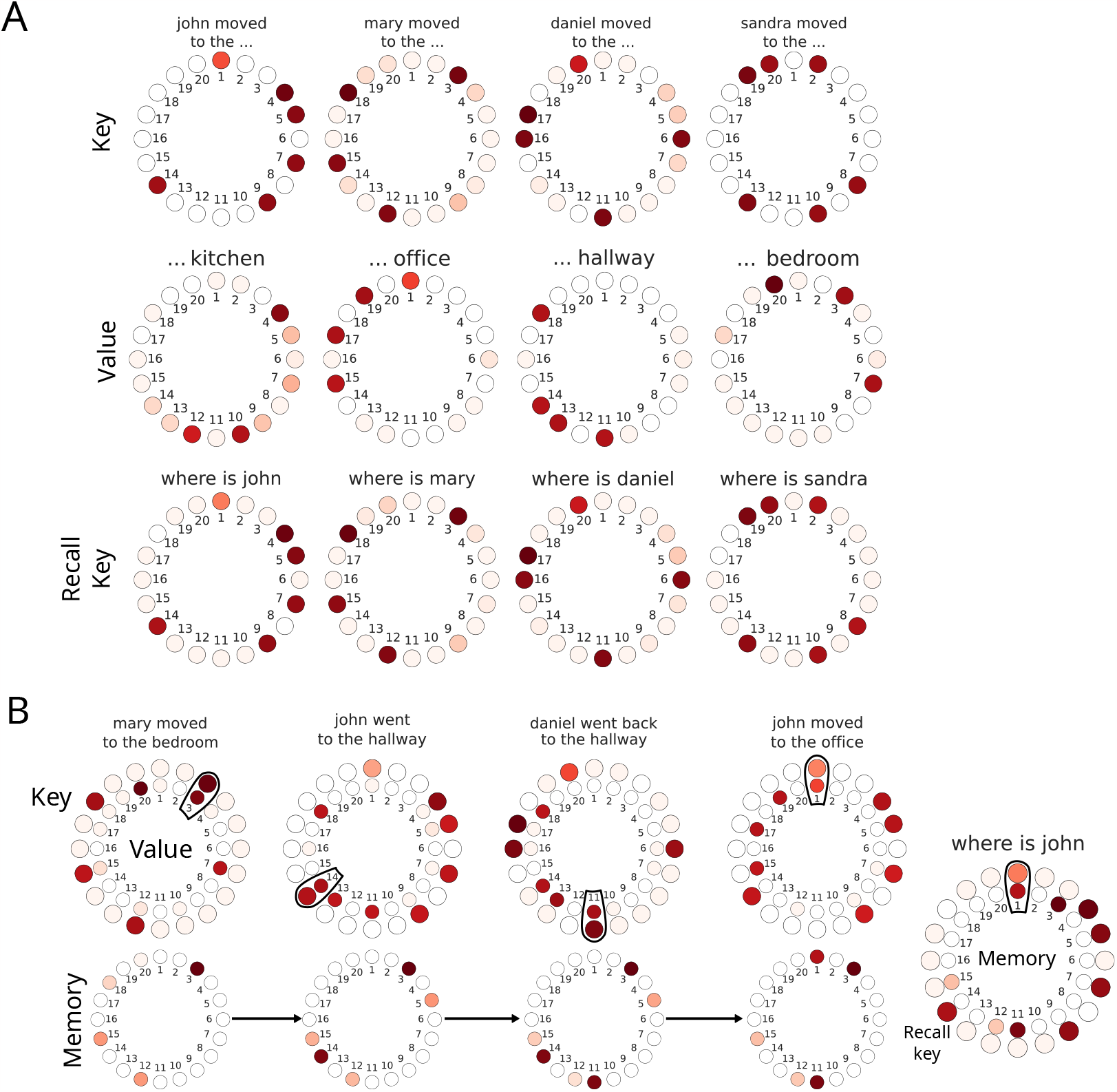
Network analysis for the Single Supporting Fact task. **A**) Projection of keys, values, and recall keys based on a non-negative matrix factorization of memory traces after network training. Keys are shown for specific persons, with representations averaged over locations and verbs. The key for John clearly activates 6 components corresponding to possible locations for John. Value representations are shown for specific locations, with representations averaged over persons and verbs. **B)** Story sample along with its respective key (top, outer ring), value (top, inner ring), and memory state after memorization (bottom). Each key and value pair predominantly overlaps in a single component, which is then memorized. Additionally, the change in John’s location in the last fact is accurately updated from component 14 to 1, causing component 14 to be deactivated due to the negative term in our Oja-type Hebbian rule.

Fig. 6A (top) shows the average activity in the key layer for a given person, where the average was taken over all possible locations and action verbs. For example, the leftmost visualized vector shows sentences of the form “john moved to the …”, where the location is marginalized out. When comparing the representations for four persons, one observes that keys effectively discriminate between persons with close to orthogonal representations. When we performed the same analysis for a given location, averaging over persons, we did not observe such a structure (not shown). In contrast, such orthogonal representations can be observed for specific locations in the value layer when an average was taken over persons and action values (Fig. 6A, middle).

In summary, we observed that keys effectively discriminate between persons, while values indicate locations. We thus define each person’s key vector as the average of the key vector projections from all possible variations of sentences for this person. Similarly, we define a location’s value vector as the average of the value vector projections from all possible variations of sentences for this location. While key- and value-representations are orthogonal to those of the same layer, we found a systematic overlap between key representations with value representations. For instance, John’s key vector primarily overlaps with specific locations indicated by the value vectors (components 1, 4, 7, and 14 overlap office, kitchen, bedroom, and hallway respectively, Fig. 6A, compare top and middle). This analysis can be similarly conducted based on the value vectors. For example, the most activated components of the office’s value overlap primarily with specific people indicated by the keys (component 1, 15, 17, and 19 overlap with John, Mary, Daniel, and Sandra respectively). The bottom row of Fig. 6A shows recall keys during queries for the location of persons (e.g., “where is john”). We observe that these recall keys are very similar to keys during storage operations for facts that include the same person.

This representation, that has been learned through local synaptic plasticity in our model, can be used to store relevant information in the trunk strengths of the pyramidal neurons of the memory layer. This is illustrated in Fig. 6 **B** where we illustrate the processing of a simple story. In the first sentence “mary moved to the bedroom”, the overlap between the key- and value-representations potentiates the trunk strength of the corresponding pyramidal neurons (component 3 in our projection), which can be observed when projecting the memory state **h**^*t*^ after the Hebbian update into the low-dimensional space (leftmost bottom representation in Fig. 6 **B**). This potentiated state is retained after the next two sentences, and new memories are added according to the presented facts. At the last presented fact “john moved to the office”, John changes his location from the hallway to the office. This change is accurately recorded in the memory: the overlapping of component 1 in the key and value represents the new location, while the deactivation of components 14, corresponding to the previous location, occurs due to the negative term in the Hebbian rule. The final state of the memory is then combined with a specific recall key from the question, “Where is John?” The answer is determined by the overlapping between their activated components, effectively corresponding to the newly activated component 1, which is part of the office representation – John’s last change of location. Hence, the readout can easily determine office as the correct answer.

In summary, the model has learned assembly representations for entities. These representations are partly orthogonal and partly overlapping. An overlap defines a potential association that can be stored in the neurons of the memory layer.

Experimental studies in humans have found clear evidence for assembly representations of celebrities and popular places in the medial temporal lobe of humans with partial overlap [38, 39]. According to our model, overlapping assemblies emerge through learning because they are needed for the storage of associations in the memory layer.

### A model for contextual fear conditioning

Contextual fear conditioning (CFC) is a classical experimental procedure to study memory in animals. In this paradigm, typically an auditory tone is paired with an aversive stimulus. After several such pairings, a freezing response can be observed when the animal is confronted with the tone alone (Fig. 7A). The CFC paradigm was instrumental in memory engram studies. Using immediate early gene cell labeling in combination with optogenetics, several studies were able to reactivate memory engram cells that were active during CFC to elicit a freezing response in the absence of the tone stimulus [2]. Interestingly, such engram reactivation even elicited the conditioned response in an amnesic condition, when synaptic consolidation was blocked after the pairings such that the tone itself did not lead to freezing [15]. Several authors hypothesized that at least in an initial stage, non-synaptic plasticity may play an important role in this learning paradigm [4, 6, 7, 10].

**Fig 7.**
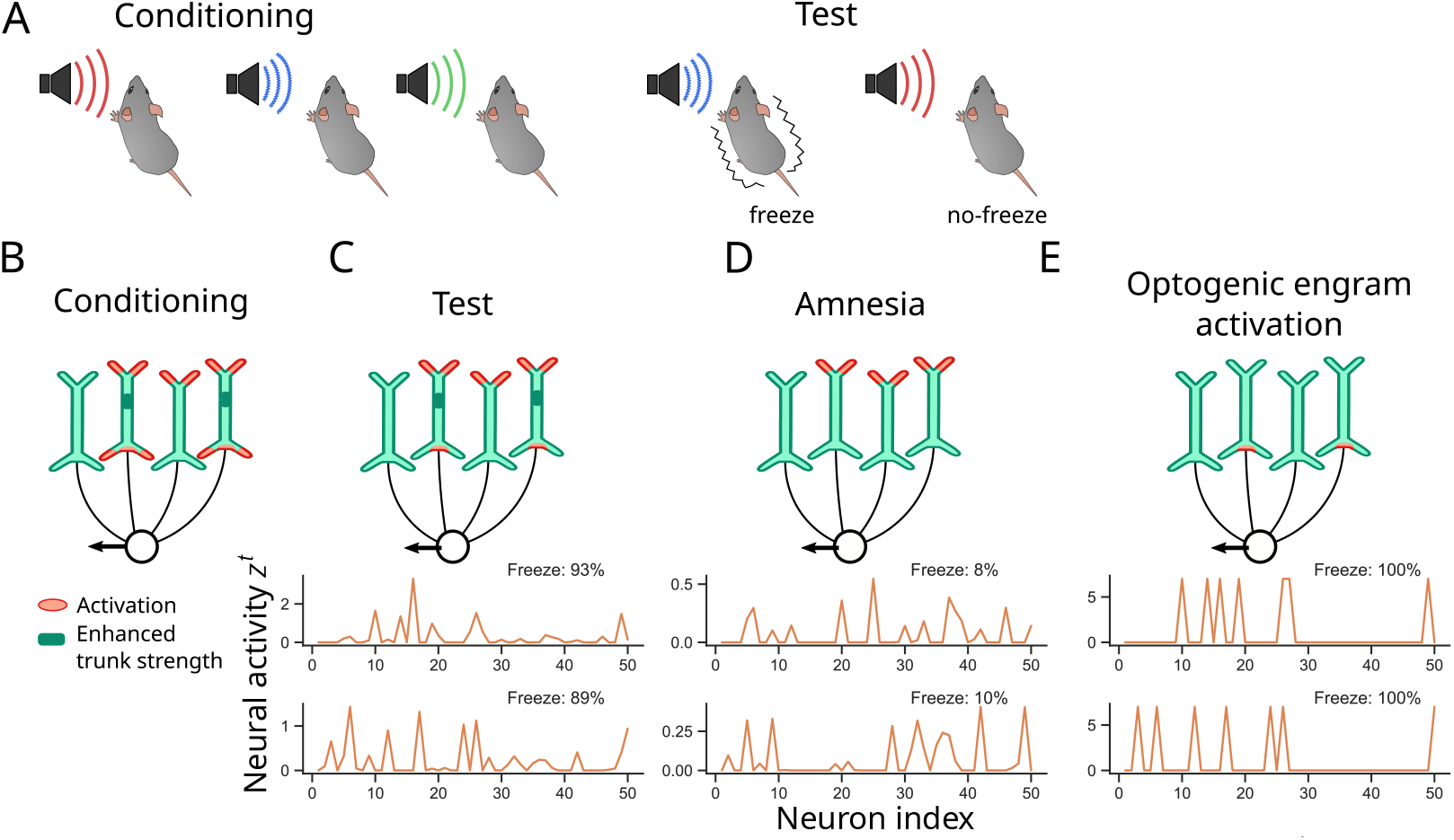
Fear conditioning and engram re-activation after amnesia. **A**) Fear conditioning paradigm. An animal is hearing several tones, where one tone (blue) is paired with a shock. After several conditioning episodes, the shock-associated stimulus is enough to elicit a freezing response. **B-E)** Schematic illustration of model behavior (top) and two example activation patterns of the 50 neurons in the memory layer in the network simulation (bottom). The percentage in the inset (“Freeze”) gives the probability that the network output assigns to a freezing response in this example. **B)** During conditioning, the patterns activate different basal and apical compartments of the memory network, enhancing the dependent trunk strength. **C)** After conditioning, the shock-associated pattern activates corresponding apical dendrites. Due to increased trunk strength, this elicits a fear response in the down-stream readout (circular neuron). Strongly active neurons in the memory layer are considered engram neurons (bottom). **D)** We set the 7 most strongly potentiated dendritic trunk strengths to zero to model the amnesic condition. Neuron responses in the memory layer are thus weak (bottom) and the fear response vanishes. **E)** Optogenetic engram stimulation re-activates engram neurons, which recovers the freeze response in the downstream readout.

We wondered whether trunk strength plasticity could provide a model for this hypothesis while being consistent with the findings of freeze responses under retrograde amnesia. To this end, we trained a network consisting of 50 neurons in the memory layer and a readout indicating a freeze response. As input to the network, we presented 50-dimensional randomly drawn sparse activation patterns modeling upstream representations of tone stimuli (see *Supplement* for details). Another input indicated the shock stimulus. Each training episode consisted of a sequence of up to 11 tone patterns (the number of patterns was sampled uniformly from {2, …, 11 }), where one of them was accompanied by a shock input. Then, two of these patterns (one shock and one non-shock) were again presented to the network. It was trained to produce a freeze response for the shock-paired input pattern. After training, the network was able to quickly associate an arbitrary input pattern to a freeze response. Tested on patterns not seen during training, it achieved 87.8% correct freeze responses and 99.5% correct non-freeze responses (over 1000 test episodes). Mechanistically, neurons in the memory layer with co-active basal and apical compartments increase their trunk strength (Fig. 7B), establishing a memory engram. When a similar pattern of apical activity is applied in a recall, the engram is activated (Fig. 7C). We found that in our model, this engram was rather sparse (Fig. 7C, bottom). The engram activates the downstream readout modeling the initiation of a freeze response.

We propose the following hypothesis for optogenetically induced freeze responses under amnesia. Here, the assumption is that the trunk potentiation is transient. It was proposed that such transient non-synaptic plasticity acts as permissive signal for LTP of recurrent connections [4] stabilizing the engram (as we do not have recurrent connections in the memory layer, this was not modeled here). However, when synaptic consolidation is blocked, recurrent connections are not altered while the trunk strengths decay. We modeled this decay of trunk strengths by setting the seven most strongly potentiated trunk strengths to zero. When the shock-associated stimulus was shown to the model afterward, the memory engram was not activated and no freeze response was elicited by the downstream readout (16.3% correct fear responses in 1000 episodes; Fig. 7D). Optogenetic reactivation was modeled by artificially activating the seven most active neurons in the original engram (i.e., setting their output to the maximum observed output over all patterns, Fig. 7E). In our model, this increased freeze response in the downstream readout to 92.5%. In summary, the hypothesis according to our model is that (a) CFC engrams are initially stored by non-synaptic plasticity mechanisms (b) the downstream readout is able to distinguish shock-paired patterns from neutral patterns, and (c) during amnesia, optogenetic reactivation of a shock-paired pattern is recognized by the readout and the fear-response is initiated. Note that the slow synaptic plasticity plays a crucial role in this model. The weight matrices have been trained previously such that a stimulus-shock association can rapidly be stored, and the downstream readout can recognize a fear memory. Hence a fear-memory engram can be recognized in the absence of further synaptic plasticity.

## Discussion

We propose in this article that memory-dependent neural processing is jointly shaped by non-synaptic and synaptic plasticity. The interaction between non-synaptic intrinsic and synaptic plasticity has been studied in previous work. These works however considered intrinsic plasticity as a homeostatic mechanism to maximize information transfer [40, 41] or to support unsupervised learning [12, 13]. The cooperation of intrinsic and synaptic plasticity has also been studied for information theoretic learning in artificial neural networks [42]. Another aspect of intrinsic plasticity is that it can regulate the dynamical behavior of recurrent network models [43–45]. In contrast to these works, our model shows that non-synaptic plasticity can be used as a memory buffer that is utilized by synaptic plasticity processes. Local plasticity for two-compartment neurons has been studied in [46], but in the context of representation learning.

The idea that fast synaptic plasticity can underlie working memory has been proposed already in [47], see [48] for a recent review. This idea has been extensively used in memory-augmented neural networks [18, 19, 25, 26, 49–51]. Here, however, synaptic plasticity was used as a fast memory buffer and networks were trained with backpropagation through time. Fast synaptic Hebbian plasticity was combined with reward-based learning in [52] to learn a navigation task. From the neuroscience perspective, non-synaptic fast plasticity is consistent with the observation that non-synaptic plasticity can act on a fast timescale [6, 9, 10]. But what could be functional advantages of non-synaptic neuron-specific plasticity? Our study indicates one potential advantage: local learning rules for synaptic connections can be used to shape the circuitry around the memory units.

The proposed network model could be improved in several directions. While we were able to show that local learning works very well when compared with BPTT (see e.g. Tab. 1), the simple memory architecture is limiting. Hence, when compared to H-Mem [19], a brain-inspired memory network without local learning, it fails on a number of bAbI tasks which can be solved by H-Mem. We were able to derive local learning rules for our model, but we failed to do so for fast memory in synaptic weight matrices. It is an interesting question whether biologically plausible learning mechanisms could be derived for this case as well. Since Hebbian branch plasticity similar to eq. (3) was found in oblique dendrites of pyramidal cells, it would be interesting to investigate models where each oblique branch acts as a memory unit similar to our 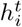. This could significantly boost the memory capacity of the model.

Another limitation of our model is that learning can take long. In the pattern association task (Fig. 3), learning of associations of two pattern-pairs needs a few hundred episodes, which increases with the number of associated pairs (Fig. 3B). Here, curriculum setups and more specialized circuit architectures are potential candidates for improvement. We emphasize however that associations can be memorized instantly after training by our model.

In summary, we have shown that non-synaptic plasticity enables memory-dependent learning with local synaptic plasticity rules. The resulting learning architecture is quite powerful, leading to results comparable to BPTT on bAbI. To the best of our knowledge, our model is the first network that is able to learn bAbI tasks with local learning rules. The involvement of non-synaptic plasticity in memory formation has been demonstrated experimentally. Our model proposes a functional role for it in synergy with synaptic plasticity.

## Materials and Methods

In this section, we provide detailed information about the model and derive the local synaptic plasticity rules.

### Memory network model

Our network model consists of 2*d* input neurons separated into apical and basal populations with activities **x**^a,*t*^ and, **x**^b,*t*^ respectively. The input neurons project to apical and basal compartments of *m* memory neurons in a fully-connected manner via weight matrices *W* ^apical^ and, *W* ^basal^ respectively. For the *i*-th neuron in this memory layer, the resulting basal activation 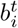 and apical activation 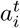 are thus given by:

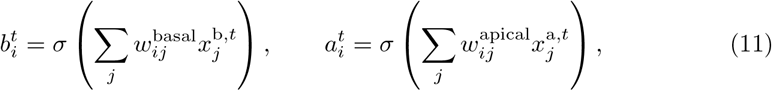

with **x**^a,*t*^ and **x**^b,*t*^ as apical and basal input vectors respectively, at time step *t* and ReLU non-linearity *σ*, defined by *σ*(*s*) = max 0, *s* . The output 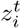 of each cell is then given by a linear combination of these two activations, whereof 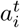 is multiplied with the trunk excitability 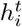:

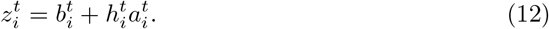

The trunk excitability 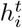 is updated according to the Hebbian update

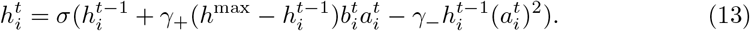

for parameters *γ*_+_, *γ*_*−*_ *>* 0.

The network output was then computed as

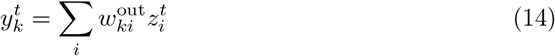

for output weight matrix *W* ^out^, where ***z***^*t*^ is determined by (12). Note, that in the reinforcement learning case, we further split this readout weight matrix into ***w***_*V*_ and *W*_*π*_, respectively, the value and policy weights, but here this separation is omitted to simplify notation. For these cases, *W* ^out^ can be interpreted as a concatenation of ***w***_*V*_ and *W*_*π*_ into a single matrix.

We distinguish between a *memorization event*, where the memory is updated according to Equation (13) and the model output is discarded, and a *memory recall event*, where the model produces an output **z**^*t*^ based on the memory state **h**^*t*^ as in Equation (12), but no memory update is performed. During memory recall events, only the apical input population is active, the basal input population is defined to be silent, i.e. **x**^b,*t*^ = **0**. In the reinforcement learning setup, we execute a memorization event followed by a memory recall event at every single time step. The network output of the recall event is then used to update the state value estimator. In the supervised learning setup, network outputs are only needed when an action is demanded (we call this a *query step* and the input at this step a *query*). Therefore, we perform memory recall events at query steps and memorization events are performed for other time steps.

### Derivation of local synaptic plasticity rules

Here, we derive gradient-based local plasticity rules for the slow synaptic weights *W* ^out^ in the output layer and *W* ^apical^, *W* ^basal^ in the memory layer (in contrast to the fast trunk strengths in the memory layer neurons).

In vanilla backpropagation through time (as commonly used for training recurrent networks), symmetrical feedback connections are used to back-propagate the error gradients through the unrolled network. These symmetrical feedback weights are generally considered biologically implausible due to the weight transport problem [53]. In this work we use the e-prop framework [28] to derive local learning rules for every synapse, that require neither the application of symmetric weights, nor the propagation of errors to preceding time steps.

#### Output layer

For the readout layer weights *W* ^out^, local learning signals are already available because the output of this layer is directly used to obtain the error signal *E*. We can therefore update output weight wout 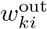 from memory layer neuron i to output neuron *k* with

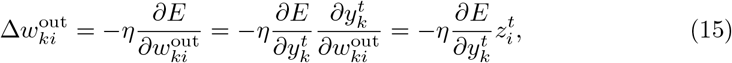

where *η >* 0 denotes the learning rate.

#### Basic e-prop framework and feedback alignment

To perform gradient descent on weights in the memory layer, we need to compute the gradients

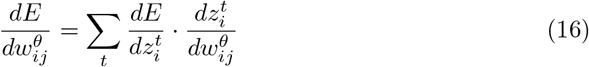

with *θ ∈* {apical, basal}. The second term 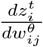 can be expressed as an eligibility trace, see below. The first term 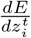 however necessitates the backpropagation of error signals through time. One core idea of the e-prop formalism [28] is to replace this gradient by a temporally local approximation,

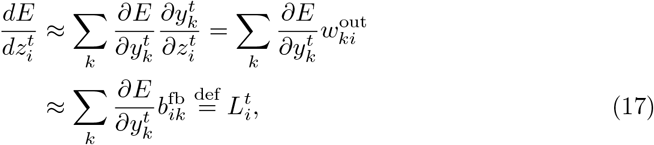

where index *k* denotes the *k*-th neuron in the output layer. Here, 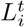 is interpreted as a neuron-specific learning signal for neuron *i* in the memory layer.

In the above equations, we distinguish between the total derivative 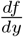 and the partial derivative 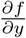. The function *f* may depend on many variables, where many of them may also depend on *y*. The total derivative takes all these dependencies into account, while in the partial derivative there is only a direct dependency of *f* on *y*. More details about this notation can be found in [28], where the e-prop formalism was proposed.

The approximation in the last line of the above equation implements feedback alignment, where the symmetric feedback weights are replaced by randomly chosen weights [54]. In our simulations, we used the adaptive e-prop method [28], where the feedback weights are initialized randomly and then undergo the same weight updates as weights in *W* ^out^: 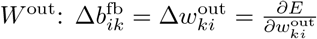 . In addition, these two weight matrices are subject to L2 weight decay. This ensures, that after some iterations *W* ^out^ and *B*^fb^ will converge to similar values.

#### Memory layer

By making use of the learning signals *L*^*t*^ from the previous section, we can find local synaptic plasticity rules to update the synaptic weights *W* ^apical^ and *W* ^basal^ of the apical and basal compartment respectively. We derive local gradients 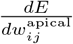 and 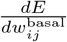 to minimize the error *E* through the e-prop formalism [28]. With these local gradients, it is not necessary to back-propagate the error signal through all time steps of the computation. Instead, eligibility traces are forward-propagated and used in conjunction with learning signals to determine the gradient required for the weight updates.

#### Apical compartment

To find the derivative 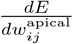 of the error function *E* with respect to apical weight 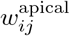, we factorize

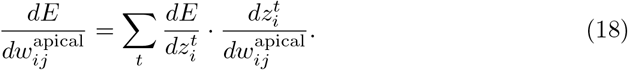

Now we use our learning signal 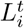 to approximate 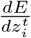, which is a core idea in the e-prop formalism [28]. We also expand 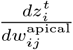 by applying the product rule, resulting in

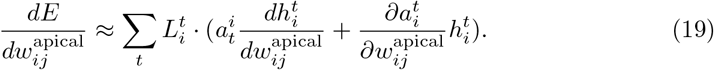

Unrolling 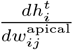 in time, we obtain

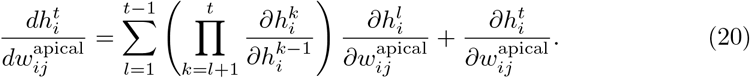

We can re-write this definition as recursive function and hereby introduce our eligibility trace 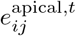:

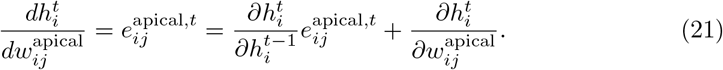

We observe, that this recurrent definition only depends on partial derivatives (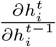 and 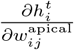) that are local in time and can therefore be easily derived from Equation (13) as:

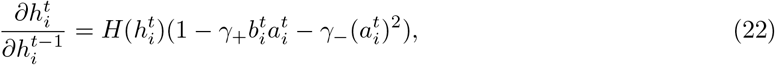

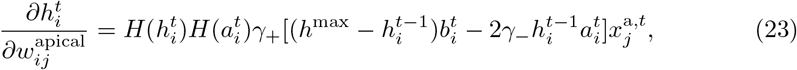

where *H*(*x*) is the Heaviside step function used as derivative of the ReLU function.

By plugging Equations (22) and (23) into (21), we obtain:

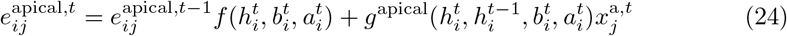

With

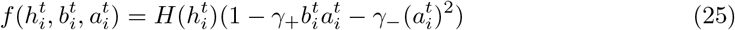

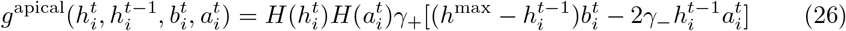

as stated in the main text. We finally use this eligibility trace to substitute 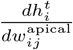 in Equation (19):

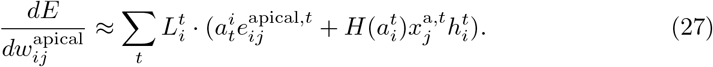

where the second term is the derivative 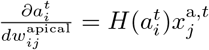 of our apical activation 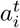 with respect to the apical weight 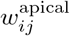.

#### Basal compartment

Analogously, we compute the derivative of the error *E* with respect to our basal weight matrix *W* ^basal^. We again start off by inserting our learning signal 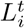:

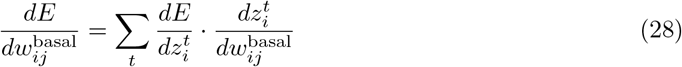

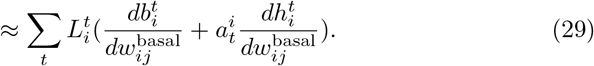

Because of our assumptions, *L*^*t*^ is only non-zero during recall events and the basal compartment does not receive any inputs at this point (i.e. 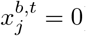), then 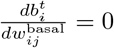.

Analogous to Equation (20), we can also express 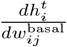 recurrently via eligibility trace 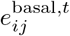.

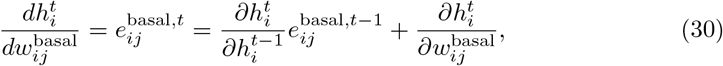

where the local derivative 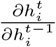 was already defined in Equation (22). To obtain 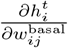 we can simply compute the derivative of Equation (13) w.r.t. 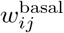:

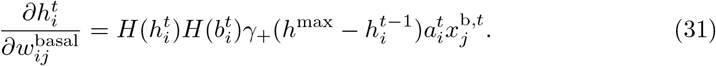

Inserting this back into (30) yields:

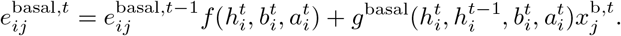

with 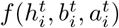 equivalent to Equation (25) and

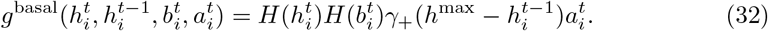

Taken together, our approximated derivative reads:

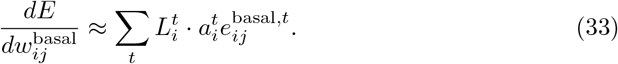

### Weight updates in simulations

The above derived approximate gradients were accumulated over mini-batches of training examples and used for parameter updates in our model. In practice, we used these approximate gradients in the Adam [55] algorithm, which amounts to a synapse-specific learning rate based on global learning rate *λ*_*Adam*_ along with a momentum update. We used the default values for other hyper-parameters of Adam which are *β*_1,*Adam*_ = 0.9, *β*_2,*Adam*_ = 0.999 and *ϵ*_*Adam*_ = 1*e −* 07. We also used an L2 regularization term for each of the synaptic weight matrices. The hyperparameters for learning are specified in the *Supplement*.

### Temporal normalization of neuron activity

In order to stabilize training, activity normalization techniques such as batch normalization and layer normalization are often used in artificial neural networks. In biological networks, such normalization may be carried out through inhibitory networks. In contrast to standard techniques, we have to ensure in our model that no information can be inferred from the future, therefore normalization needs to be performed in an online manner. We normalize apical and basal activity vectors ***a***^*t*^ and ***b***^*t*^ component-wise over the temporal dimension *t*. In our case, we use a variance-mean online computation based on [56]:

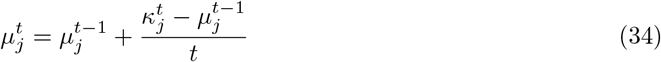

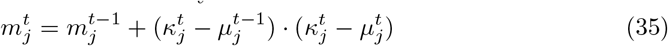

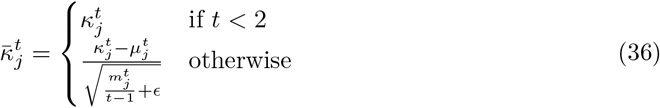

with ***κ***^*t*^ ∈ {***a***^*t*^, ***b***^*t*^} at time *t*, online approximated mean ***μ***^*t*^ and cumulative squared difference ***m***^*t*^ between input and mean. We further define ***μ***^0^ = ***m***^0^ = **0**.

For each task, we calculated ***μ***^*t*^ and ***m***^*t*^ online during the training phase across all training samples to obtain ***μ***_train_ and ***m***_train_ to apply them as constants for the normalization during inference. This procedure is a common practice and coherent with standard normalization techniques like Batch Normalization [57].

### Code source

Source code under GNU General Public License v3.0 is available at https://github.com/igiTUGraz/tsp.

## References

1. Semon RW. The mneme. G. Allen & Unwin Limited; 1921.

2. Tonegawa S, Pignatelli M, Roy DS, Ryan TJ. Memory engram storage and retrieval. Current opinion in neurobiology. 2015;35:101–109.

3. Josselyn SA, Tonegawa S. Memory engrams: Recalling the past and imagining the future. Science. 2020;367(6473):eaaw4325.

4. Mozzachiodi R, Byrne JH. More than synaptic plasticity: role of nonsynaptic plasticity in learning and memory. Trends in neurosciences. 2010;33(1):17–26.

5. Pignatelli M, Ryan TJ, Roy DS, Lovett C, Smith LM, Muralidhar S, et al. Engram cell excitability state determines the efficacy of memory retrieval. Neuron. 2019;101(2):274–284.

6. Titley HK, Brunel N, Hansel C. Toward a neurocentric view of learning. Neuron. 2017;95(1):19–32.

7. Jacob AD, Mocle AJ, Frankland PW, Josselyn SA. Neuronal Excitability in Memory Allocation: Mechanisms and Consequences. In: The Oxford Handbook of the Neurobiology of Learning and Memory;.

8. Zhang W, Linden DJ. The other side of the engram: experience-driven changes in neuronal intrinsic excitability. Nature Reviews Neuroscience. 2003;4(11):885–900.

9. Gallistel CR, Matzel LD. The neuroscience of learning: beyond the Hebbian synapse. Annual review of psychology. 2013;64:169–200.

10. Gallistel CR, Balsam PD. Time to rethink the neural mechanisms of learning and memory. Neurobiology of learning and memory. 2014;108:136–144.

11. Foldiak P. Forming sparse representations by local anti-Hebbian learning. Biological cybernetics. 1990;64(2):165–170.

12. Triesch J. Synergies between intrinsic and synaptic plasticity in individual model neurons. Advances in neural information processing systems. 2004;17.

13. Savin C, Joshi P, Triesch J. Independent component analysis in spiking neurons. PLoS computational biology. 2010;6(4):e1000757.

14. Weston J, Bordes A, Chopra S, Mikolov T. Towards AI-Complete Question Answering: A Set of Prerequisite Toy Tasks. In: Bengio Y, LeCun Y, editors. 4th International Conference on Learning Representations, ICLR 2016, San Juan, Puerto Rico, May 2-4, 2016, Conference Track Proceedings; 2016.

15. Ryan TJ, Roy DS, Pignatelli M, Arons A, Tonegawa S. Engram cells retain memory under retrograde amnesia. Science. 2015;348(6238):1007–1013.

16. Larkum ME, Nevian T, Sandler M, Polsky A, Schiller J. Synaptic integration in tuft dendrites of layer 5 pyramidal neurons: a new unifying principle. Science. 2009;325(5941):756–760.

17. Körding KP, König P. Learning with two sites of synaptic integration. Network: Computation in neural systems. 2000;11(1):25–39.

18. Sukhbaatar S, Szlam A, Weston J, Fergus R. End-To-End Memory Networks. In: Cortes C, Lawrence N, Lee D, Sugiyama M, Garnett R, editors. Advances in Neural Information Processing Systems. vol. 28. Curran Associates, Inc.; 2015.

19. Limbacher T, Legenstein R. H-Mem: Harnessing synaptic plasticity with Hebbian Memory Networks. In: Larochelle H, Ranzato M, Hadsell R, Balcan MF, Lin H, editors. Advances in Neural Information Processing Systems. vol. 33. Curran Associates, Inc.; 2020. p. 21627–21637.

20. Magee JC, Johnston D. Plasticity of dendritic function. Current opinion in neurobiology. 2005;15(3):334–342.

21. Debanne D, Poo MM. Spike-timing dependent plasticity beyond synapse-pre-and post-synaptic plasticity of intrinsic neuronal excitability. Frontiers in synaptic neuroscience. 2010;2:21.

22. Debanne D, Inglebert Y, Russier M. Plasticity of intrinsic neuronal excitability. Current opinion in neurobiology. 2019;54:73–82.

23. Losonczy A, Makara JK, Magee JC. Compartmentalized dendritic plasticity and input feature storage in neurons. Nature. 2008;452(7186):436–441.

24. Oja E. Simplified neuron model as a principal component analyzer. Journal of mathematical biology. 1982;15(3):267–273.

25. Schmidhuber J. Learning to control fast-weight memories: An alternative to dynamic recurrent networks. Neural Computation. 1992;4(1):131–139.

26. Ba J, Hinton GE, Mnih V, Leibo JZ, Ionescu C. Using Fast Weights to Attend to the Recent Past. In: Lee D, Sugiyama M, Luxburg U, Guyon I, Garnett R, editors. Advances in Neural Information Processing Systems. vol. 29. Curran Associates, Inc.; 2016. p. 4331–4339.

27. Lillicrap TP, Santoro A. Backpropagation through time and the brain. Current Opinion in Neurobiology. 2019;55:82–89.

28. Bellec G, Scherr F, Subramoney A, Hajek E, Salaj D, Legenstein R, et al. A solution to the learning dilemma for recurrent networks of spiking neurons. Nature communications. 2020;11(1):1–15.

29. Lake BM, Salakhutdinov R, Tenenbaum JB. Human-level concept learning through probabilistic program induction. Science. 2015;350(6266):1332–1338. doi:10.1126/science.aab3050.

30. Snell J, Swersky K, Zemel RS. Prototypical Networks for Few-shot Learning; 2017.

31. Schulman J, Wolski F, Dhariwal P, Radford A, Klimov O. Proximal policy optimization algorithms. arXiv preprint arXiv:170706347. 2017;.

32. Chudasama Y. In: Stolerman IP, editor. Delayed (Non)Match-to-Sample Task. Berlin, Heidelberg: Springer Berlin Heidelberg; 2010. p. 372–372.

33. van der Maaten L, Hinton G. Visualizing Data using t-SNE. Journal of Machine Learning Research. 2008;9(86):2579–2605.

34. Hochreiter S, Schmidhuber J. Long short-term memory. Neural computation. 1997;9(8):1735–1780.

35. Olton DS, Collison C, Werz MA. Spatial memory and radial arm maze performance of rats. Learning and motivation. 1977;8(3):289–314.

36. He K, Zhang X, Ren S, Sun J. Delving deep into rectifiers: Surpassing human-level performance on imagenet classification. In: Proceedings of the IEEE international conference on computer vision; 2015. p. 1026–1034.

37. Lee D, Seung HS. Algorithms for non-negative matrix factorization. Advances in neural information processing systems. 2000;13.

38. Quiroga RQ. Neuronal codes for visual perception and memory. Neuropsychologia. 2016;83:227–241.

39. Ison MJ, Quiroga RQ, Fried I. Rapid encoding of new memories by individual neurons in the human brain. Neuron. 2015;87(1):220–230.

40. Laughlin S. A simple coding procedure enhances a neuron’s information capacity. Zeitschrift für Naturforschung c. 1981;36(9-10):910–912.

41. Stemmler M, Koch C. How voltage-dependent conductances can adapt to maximize the information encoded by neuronal firing rate. Nature neuroscience. 1999;2(6):521–527.

42. Li Y, Li C. Synergies between intrinsic and synaptic plasticity based on information theoretic learning. PloS one. 2013;8(5):e62894.

43. Schrauwen B, Wardermann M, Verstraeten D, Steil JJ, Stroobandt D. Improving reservoirs using intrinsic plasticity. Neurocomputing. 2008;71(7-9):1159–1171.

44. Steil JJ. Online reservoir adaptation by intrinsic plasticity for backpropagation–decorrelation and echo state learning. Neural networks. 2007;20(3):353–364.

45. Lazar A, Pipa G, Triesch J. Fading memory and time series prediction in recurrent networks with different forms of plasticity. Neural Networks. 2007;20(3):312–322.

46. Bredenberg C, Lyo B, Simoncelli E, Savin C. Impression learning: Online representation learning with synaptic plasticity. Advances in Neural Information Processing Systems. 2021;34:11717–11729.

47. Sandberg A, Tegnér J, Lansner A. A working memory model based on fast Hebbian learning. Network: Computation in Neural Systems. 2003;14(4):789.

48. Lansner A, Fiebig F, Herman P. Hebbian fast plasticity and working memory. arXiv preprint arXiv:230406626. 2023;.

49. Miconi T, Stanley K, Clune J. Differentiable plasticity: training plastic neural networks with backpropagation. In: International Conference on Machine Learning. PMLR; 2018. p. 3559–3568.

50. Miconi T, Kay K. An active neural mechanism for relational learning and fast knowledge reassembly. bioRxiv. 2023; p. 2023–07.

51. Rohani SRR, Hedayatian S, Baghshah MS. BIMRL: Brain Inspired Meta Reinforcement Learning. In: 2022 IEEE/RSJ International Conference on Intelligent Robots and Systems (IROS). IEEE; 2022. p. 9048–9053.

52. Kumar MG, Tan C, Libedinsky C, Yen SC, Tan AYY. One-shot learning of paired associations by a reservoir computing model with Hebbian plasticity. arXiv preprint arXiv:210603580. 2021;.

53. Crick F. The recent excitement about neural networks. Nature. 1989;337(6203):129–132.

54. Lillicrap T, Cownden D, Tweed D, Akerman C. Random synaptic feedback weights support error backpropagation for deep learning. Nature Communications. 2016;7:13276. doi:10.1038/ncomms13276.

55. Kingma DP, Ba J. Adam: A Method for Stochastic Optimization. arXiv preprint arXiv:14126980. 2014;doi:10.48550/ARXIV.1412.6980.

56. Welford B. Note on a method for calculating corrected sums of squares and products. Technometrics. 1962;4(3):419–420.

57. Ioffe S, Szegedy C. Batch Normalization: Accelerating Deep Network Training by Reducing Internal Covariate Shift; 2015.

